# Out of Antarctica: new insights into Antarctic Subcluster 5.2 picocyanobacteria based on high-quality genomes

**DOI:** 10.64898/2025.12.17.694815

**Authors:** Benoit Durieu, Valentina Savaglia, Mick Van Vlierberghe, Valérian Lupo, Denis Baurain, Annick Wilmotte, Luc Cornet

## Abstract

2.

*Synechococcus*-like cyanobacteria are cosmopolitan unicellular picocyanobacteria that have colonized diverse aquatic and terrestrial habitats. The so-called ‘subcluster 5.2’ represents a particularly diversified subgroup, including marine and freshwater organisms adapted to extreme conditions, notably polar environments. We increased the genomic representation of polar taxa in this subcluster by reconstructing new high-quality genomes from five Antarctic lacustrine strains and one Arctic freshwater isolate using a combination of small Illumina and long Nanopore reads. A maximum likelihood (ML) phylogenomic analysis of these new assemblies combined with all publicly available good quality assemblies of the subcluster 5.2 suggests evidence of a dispersal process from Antarctica. Indeed, the topology of the phylogenomic tree indicates one basal Antarctic lineage followed by the emergence of two lineages, one Antarctic and one non-Antarctic (Spain). This finding is further supported by a 16S rRNA ML phylogenetic and a pangenomic analysis. Although secondary colonization of Antarctica by cyanobacteria following the cooling of the continent 34 million years ago has been reported, this study is the first to support an ‘Out-of-Antarctica’ scenario inside subcluster 5.2.

**Impact statement:** This study provides novel insights into the evolutionary history and biogeography of *Synechococcus*-like picocyanobacteria. By expanding the genomic sampling of polar subcluster 5.2, we reveal that Antarctic taxa are not monophyletic and are intermixed with non-Antarctic lineages, suggesting an unprecedented Out-of-Antarctica dispersal scenario. Our results, supported by 16S rRNA, phylogenomic and pangenomic analyses, highlight the role of polar regions as both a refuge and a source of microbial diversity. This work advances our understanding of cyanobacterial adaptation, radiation and genome evolution in extreme environments.

**Data summary:** Raw sequencing reads and genome assemblies from this study have been deposited in the European Nucleotide Archive (ENA) under Project accession number PRJEB103944. Raw sequencing reads are available under accession numbers ERR15933635 to ERR15933638, ERR15933641 to ERR15933644, ERR15903357, ERR15903358, ERR15905256 and ERR15905257. The assemblies can be accessed under accession numbers GCA_977927515, GCA_977927525, GCA_977927535, GCA_977927545, GCA_977927555 and GCA_977929215. The scripts and code as well as large supplementary files generated during this study are available on Figshare and can be downloaded from https://doi.org/10.6084/m9.figshare.30772781 [1]. Supplementary data files are also provided and contains supplementary figures (Fig. S1-S4) and tables (Table S1-S6).

## 5. Introduction

*Synechococcus*-like cyanobacteria are a group of picocyanobacteria, including notably the *Synechoccocus* and *Cyanobium* genera, of which taxonomy remains under debate because their similar morphology has led to classification into polyphyletic groups [2, 3]. They have colonized a wide range of aquatic and terrestrial habitats, exhibiting a cosmopolitan distribution from tropical to polar regions [4–6]. Among these lineages, subcluster 5.2 is a particularly diversified subgroup, notable for its low 16S rRNA identity [7]. Originally described as a marine group [8, 9], subsequent studies have shown that subcluster 5.2 includes also non-marine microorganisms thriving in variable and extreme conditions, encompassing a wide range of salinity and temperature [5, 7, 10–12].

Phylogenetic studies on 16S rRNA data revealed the presence of one group of Antarctic cyanobacteria within subcluster 5.2 [5, 13–15], among which a single strain sampled from the marine-derived Ace Lake (Vestfold Hills) has been studied through genomic analyses [16, 17]. This strain plays a major ecological role in the lake by blooming to high levels in a seasonal dynamic influenced by polar light cycle [18].

Cyanobacteria present in Antarctica are either cosmopolitan or endemic to the continent [19]. Multiple colonization events from other continents have been documented, from lineages scattered across the cyanobacterial phylogeny [20]. Nevertheless, no events of dispersal from Antarctica towards another location, i.e., an Out-of-Antarctica scenario, have been reported to date. In this study, we increased the representation of subcluster 5.2 with polar strains by reconstructing six high-quality genome assemblies from five Antarctica and one Arctic isolates. These new genome assemblies allow us to present a phylogenomic and pangenomic analysis demonstrating that Antarctic taxa from subcluster 5.2 do not form a monophyletic group but instead are intermixed with non-Antarctic lineages. Interestingly, those secondarily non-Antarctic cyanobacteria contain a pseudogeneized version of a gene that is likely functional in their Antarctic relatives. These findings have important evolutionary history implications, as in addition to multiple colonization events of Antarctica, our results open the possibility of a reverse, Out-of-Antarctica, scenario in this case.

## 6. Methods

### Sampling

We reconstructed the genome of four polar strains of *Cyanobium* sp. from the BCCM/ULC Cyanobacteria Collection (**Table 1**). Three strains were isolated by W. Vincent in Antarctic lakes from the Livingston Island (ULC081: Limnopolar Lake, ULC082: Chester Cone, ULC084: Laguna Chica), while strain ULC065 was also isolated by W. Vincent, but in Bylot Island (Arctic, Canada). Additionally, two Antarctic *Synechococcus* sp. strains (CS-602 and CS-603) isolated from Pendant Lake and Lake Abraxis, respectively, were ordered from the Australian National Algae Culture Collection, ANACC (**Table 1**). Strains were grown in BG11 medium at a temperature of 12 °C.

**Table 1.** Culture collection, isolation sources and ENA accessions of the strains sequenced in the present study.

| Strain | Culture collection | Isolation source | Deposit year | ENA accession |
| --- | --- | --- | --- | --- |
| ULC065 | BCCM/ULC | Bylot Island, Canada, Arctic | 2011 | GCA_977927515.1 |
| ULC081 | BCCM/ULC | Limnopolar Lake, Antarctica | 2006 | GCA_977927525.1 |
| ULC082 | BCCM/ULC | Chester Cone, Antarctica | 2011 | GCA_977927535.1 |
| ULC084 | BCCM/ULC | Laguna Chica, Antarctica | 2011 | GCA_977927545.1 |
| CS-602 | ANACC | Pendant Lake, Antarctica | 1993 | GCA_977927555.1 |
| CS-603 | ANACC | Lake Abraxis, Antarctica | 1993 | GCA_977929215.1 |

### DNA extraction and de novo sequencing

For each strain, 6 ml of liquid culture was centrifuged at 12,000 g to remove water and obtain biomass pellets for DNA extraction using the GenElute™ Bacterial Genomic DNA Kit (Sigma-Aldrich, St. Louis, MO). DNA amount and quality were checked on electrophoresis gel (0,8 % agar, 80 V, 100 mins) and with the Quant-iT™ PicoGreen™ dsDNA Assay Kit (Thermo Fisher Scientific, Waltham, MA).

Short paired-end read sequencing was performed for the six strains on an Illumina MiSeq 600 cy V3 sequencer (2 x 300 bp) and the Nextera XT DNA Library preparation Kit (Illumina, San Diego, CA, USA). Long read sequencing was performed on a MinION Flow Cell R9.4.1 (Oxford Nanopore Technologies, UK) after barcoding and ligation sequencing preparation using Native Barcoding Kit 24 (Oxford Nanopore Technologies, UK). A sequencing depth of 100 x was expected for the two sequencing technologies, based on a hypothesized average genome size of 3 Mbp.

### Genome assembly and quality assessment

Raw reads from Illumina sequencing were first filtered with PRINSEQ [21] with custom parameters (-min_len: 80, -min_qual_mean: 15, -trim_qual_right: 25, -qual_noscale) and then with Trimmomatic [22] to remove Illumina adaptors (leading: 20, trailing: 20, slindingwindow: 10:20, crop: 270, minlen: 80). Base-calling of the MinION raw reads was performed using the Guppy basecaller (Oxford Nanopore Technologies, UK) and a filter step was applied with Nanofilt [23] to retain only sequences with Q-score ≥ 10. Hybrid de novo assembly of filtered reads was then performed with Unicycler (-- no_correct, --mode normal; Wick et al., 2017). We identified cyanobacterial contigs in the hybrid assemblies using Kraken2 [25] default parameters and databases, and those cyanobacterial contigs were merged after checking for redundancies using mummer and mummerplot [26]. CheckM [27] was then used with the “Taxonomic and Lineage-specific” workflow to determine completeness and contamination. Information about length, taxonomy and mummer redundancies of the contigs is provided in **Table S1**.

### Multilocus maximum likelihood phylogeny (phylogenomic analysis)

The 2336 genomes assemblies of the order Synechococcales available as of 7 June 2024 on the NCBI Assembly database were downloaded using the Genome-downloader.nf workflow implemented in the GEN-ERA toolkit [28]. These assemblies were compared to our six polar assemblies by Average Nucleotide Identity (ANI) using the ANI.nf workflow (also from GEN-ERA), with default parameters [28]. To encompass the full diversity of subcluster 5.2, we conserved 291 assemblies sharing at least 77% of ANI with one or more of our polar assemblies, a threshold chosen slightly below the value used by Dufresne et al. [8] to discriminate subclusters. Quality criteria (contamination ≤ 5%, completeness ≥ 95%) were computed using the GENcontams.nf workflow (GEN-ERA), with default parameters [28]. When multiple assemblies were available for a single cyanobacterial strain, only the highest-quality assembly was retained for further analyses based on completeness and contamination. This results in the selection of 48 of the 291 assemblies for phylogenomic analysis. We then added *Synechocystis* PCC 7509 (GCF_000332075.2) as an outgroup (**Table S2**; metadata of the 49 selected assemblies + our six new polar ones). A phylogenetic inference was performed on these 55 genomes using the Phylogeny.nf workflow from the GEN-ERA toolbox, with default parameters [26]: 476 Orthologous groups (OGs), corresponding to unicopy genes shared by all genomes, were retrieved [28]. A multi locus maximum likelihood phylogeny using RAxML [29] with 1000 rapid bootstraps under the PROTGAMMALGF model implemented in the Phylogeny.nf workflow (GEN-ERA) [28] was performed on these 476 OGs. The phylogenomic tree was visualized with iTOL v7 [30] to delineate the different polar clusters.

### 16S rRNA phylogeny

16S rRNA genes were extracted from assemblies with RNAmmer v1.2 [31]. We then retrieved all available sequences from the NCBI “core_nt” database (carried out on 10 March 2025) at 97% similarity threshold, to encompass the diversity of the phylogenomic “cluster D”. The retained 16S rRNA gene sequences were aligned using MAFFT v8.2.12 [32] using default parameters and a phylogenetic tree was inferred using RAxML [29] with 100 rapid bootstraps under the GTRGAMMA model. Geographic locations for the whole dataset were retrieved in batch from GenBank using EFetch (Entrez Fetch) utility [33]. The final tree was visualized with iTOL v7 [30].

### Genome annotations and pangenomic analysis

Among the 55 assemblies included in the phylogenomic analysis, 44 genomes were retained for the pangenomic analysis. Selection was based on assemblies containing no more than 100 contigs and showing less than 50% heterogeneity, according to CheckM results.

Pangenomic analysis was performed using the ‘anvi-pan-genome’ program in Anvi’o, after genome formatting with ‘anvi-gen-contigs-database’ and annotation using ‘anvi-run-ncbi-cogs’, ‘anvi-run-kegg-kofams’, and ‘anvi-run-pfams’ [34–40] (Figshare identifier: 10.6084/m9.figshare. 30772781). The workflow used BLASTp to compute the amino acid identity level between all ORFs and removes matches below a bitscore threshold (minbit heuristic = 0.5; Altschul et al., 1990). Homologous gene clusters (GCs) were then determined using the Markov Clustering Algorithm (MCL inflation = 8; [35, 36]) (Figshare identifier: 10.6084/m9.figshare.30772781).

Functional annotations significantly enriched (q-value < 0.05) in Antarctic genomes compared to non-Antarctic genomes were identified with the program ‘anvi-compute-functional-enrichment-in-pan’. A false discovery rate correction was applied to p-values generated using the Rao score test for equality of proportions to account for multiple testing. To strengthen the robustness of our functional annotation analysis, we used the COG and KOfam/Pfam databases and retained only enriched functions for which the GCs corresponded across both databases.

### In-depth analysis of specific Antarctic gene clusters and functional enrichment analysis

The homologous gene clusters (GCs) inferred from the pangenomic analysis were examined with respect to their Antarctic or non-Antarctic origin. GCs specific to Antarctic (i.e., present only in Antarctic assemblies or in all Antarctic assemblies and not in their closest relatives) were counted using a custom Perl script (Figshare identifier: 10.6084/m9.figshare. 30772781). From these GCs, those with functional annotations significantly overrepresented in Antarctic assemblies (i.e., genes *InsA* and *HigB*) were selected for further analysis (Figshare identifier: 10.6084/m9.figshare.30772781). As *HigB* was sometimes unexpectedly absent from genomes despite its supposedly tight functional association with *HigA*, we applied a more sensitive detection method using the program Forty-Two [37], as in Lupo et al. [38]. To this end, *HigB* protein sequences were aligned with Clustal Omega [39], manually curated, and a PhyML tree [40] was inferred under the GTRGAMMA model to separate the protein sequences annotated as HigB in homologous groups [41]. Each homologous group of HigB protein sequences was then aligned with MAFFT [32] and searched into the 44 assemblies using the TBLASTN mode of Forty-Two. These searches allowed us to classify the protein sequences find by Forty-Two as (a) HigB (97% similarity, green), (b) potential HigB (85% similarity, grey), (c) potentially not HigB (< 74% similarity, black), (d) partial sequences (turquoise), (e) truncated sequences (brown) and (f) undetermined HigB (presence of “X”, red). The exact genomic positions of the HigB protein sequences detected with Forty-Two were then added to the GFF file generated by Anvi’o for each assembly using custom Python scripts (Figshare: 10.6084/m9.figshare.30772781). The resulting GFF files were used in GeneSpy [42] to visualize the genomic context of *HigB* gene sequences across all assemblies.

## 7. Results

### General characteristics of the assemblies

For our six new polar assemblies, the number of cyanobacterial contigs identified by Kraken2 ranges from 1 (ULC065) to 13 (CS-602) for a minimum completeness of 99.46% (UC081 and CS-602) and a maximal contamination level of 0.54% (ULC081 and ULC065). Statistics for these new assemblies and the 49 others retrieved from NCBI Assembly database are summarized in Table S2. Briefly, the maximum number of contigs is 224 (*Synechococcus lacustris* GCF_003011125.1), while the minimum completeness is 97.69% (*Cyanobium* sp. A1C AMD GCF_024346495.1) and the maximum contamination level is 1.72% (*Synechococcus lacustris* GCF_003011125.1). Genome size ranges from 2,164,299 bp (Ircinia ramosa symbiont *Synechococcus* sp. SB0662 GCA_009840445.1) to 4,074,513 bp (freshwater *Cyanobium* sp. FGCU6 GCA_024958655.1), GC content from 40.62% (*Synechococcus* sp. PCC 7502 GCF_000317085.1) to 69.59% (*Cyanobium* sp. LC18 GCF_009834675.1) and the number of genes from 2140 (*Synechococcus* sp. SB0662 GCA_009840445.1) to 4280 (*Cyanobium* sp. FGCU6 GCA_024958655.1).

### Antarctic genomes are paraphyletic in subcluster 5.2

The phylogenomic analysis reveals that Antarctic and Arctic genomes are not monophyletic at 77% of ANI, those being found in respectively two and three different ANI clusters within subcluster 5.2 (**Fig. 1**). These clusters contain genomes from microorganisms inhabiting glacial or temperate lakes, as well as marine environments. **Cluster A** encompasses freshwater lacustrine *Cyanobium*/*Synechococcus* taxa is present with a higher number of, Spain, USA), one from the post-glacial freshwater lake Constance (Switzerland), whereas strain CS-1332 is from Arctic (Nunavut, Ellesmere Island, Canada). **Cluster B** is composed of four freshwater lacustrine taxa from temperate locations (Mexico, Spain) and two from Arctic (Canada). **Cluster C** is a mixture of lacustrine (New Zealand, Spain, Argentina) and marine (Black Sea, Bulgaria, and Menai Strait, UK) taxa, while 11F2 reportedly is an Antarctic strain from Edmonson Point (Victoria Land, Antarctica). Unfortunately, we could not further analyze strain 11F2 because it was abnormally too close to the Spanish strains: 100% ANI (**Table S3**), with almost the same number of genes and genomes sizes (**Table S2**), suggesting a traceability issue that we were unable to resolve even after re-ordering and sequencing the strain anew (99.99% ANI between the assembly used in this study and the re-sequenced strain). **Cluster E** is characterized by taxa from a multitude of lacustrine sources, such as Arctic (Nunavut, Bylot Island, Canada), freshwater post-glacial alpine lakes (Lake Lugano, Switzerland, and Lake Mondsee, Austria), alkaline lake Atexcac (Mexico), artificial freshwater lake in Czech Republic (Mostecké Jezero), freshwater pond in Montréal (Canada), the freshwater Loriguilla Reservoir (Chulilla, Spain) and an undetermined source of water found in the digestive system of a honeybee from Beijing (China).

**Fig. 1.**
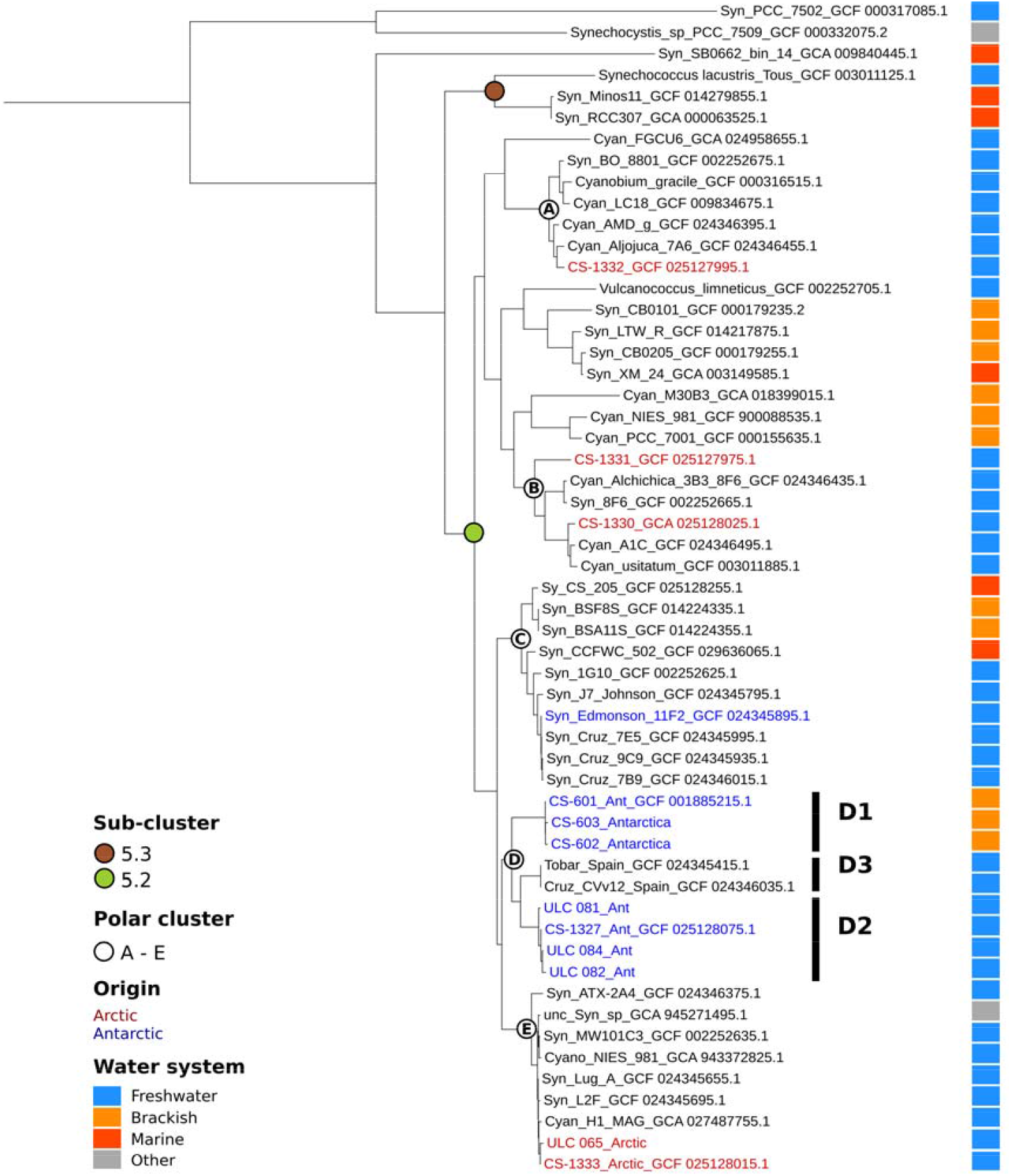
Multilocus maximum likelihood phylogeny with 1000 rapid bootstraps under the PROTGAMMALGF model of 55 cyanobacterial assemblies inferred from a supermatrix of 476 genes totalizing 86,369 unambiguously aligned amino acid positions and 0.21% of missing characters.

Regarding **Cluster D**, it encompasses freshwater and brackish Antarctic taxa, respectively from Livingston Island (near Antarctic peninsula) and from Vestfold Hills (Princess Elisabeth Land) and two freshwater taxa originating from lakes situated in the mountains of the Serrania de Cuenca (Central-Eastern Spain). Interestingly, the Antarctic taxa from cluster D are grouped in two monophyletic subclusters separated by the Spanish assemblies (**Fig. 1**). The first contains the brackish Antarctic genomes CS-601, CS-602 and CS-603 from different lakes of Vestfold Hills, which share 99.2 to 99.6 % ANI (group D1). The second includes the four freshwater Antarctic genomes ULC081, ULC082, ULC084 and CS-1327, sampled from different lakes of Livingston Island and sharing 98.1 to 99.9 % ANI (group D2) (**Table S3**). The group D2 is more genetically distant from the group D1 (81.4 to 81.6 % ANI) than from the Spanish group (group D3), comprising the strains Cruz_CV_v_12 and Tobar12_5_mg (84.6 to 84.7 % ANI).

### Enrichment of cluster D with environmental 16S rRNA sequences confirms the existence of non-Antarctic cyanobacteria falling between the two Antarctic groups

The paraphyletic nature of Antarctic taxa in cluster D was confirmed by the phylogenetic analysis of their 16S rRNA sequences. With a bootstrap above 90%, this analysis confirms that cluster D contains non-Antarctic taxa between the two Antarctic groups (**Fig. 2**). We could not recover any 16S rRNA sequence from GCF_024345415.1 (*Synechococcus* sp. Tobar12_5mg) and GCF_024346035.1 (*Synechococcus* sp. Cruz_CVv12) assemblies, but other 16S rRNA sequences were retrieved from the NCBI “core_nt” database did fall in cluster D. These non-Antarctic sequences were from polar (Greenland), alpine (Pyrenees), temperate (Portugal, Acores) and coastal marine (Russia) locations.

**Fig. 2.**
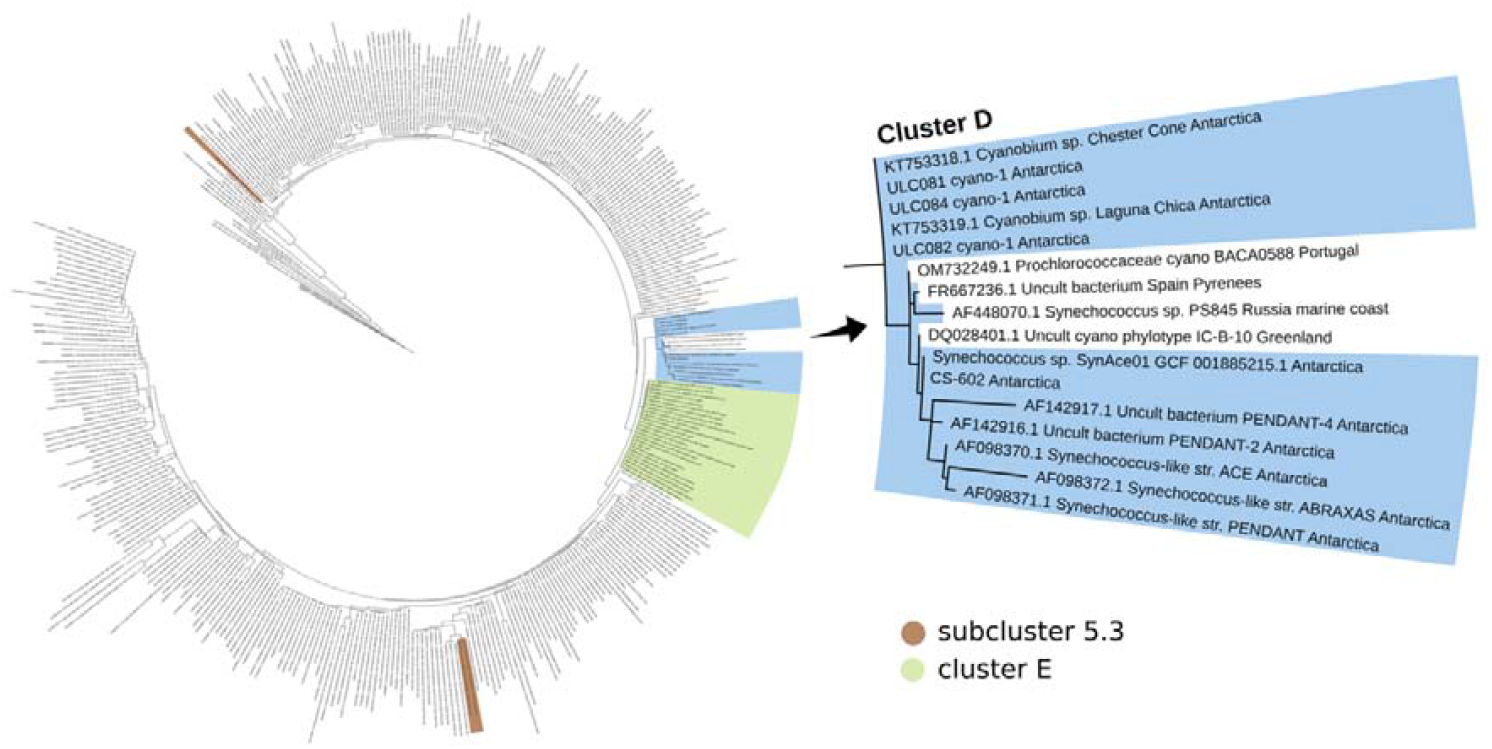
Maximum likelihood phylogeny of the 405 16S rRNA sequences contextualizing the Antarctic Cluster D, with 100 rapid bootstraps under the GTRGAMMA model for an alignment of 746 unambiguously aligned nucleotide positions and 6.63% missing characters.

We also note that subcluster 5.3 is split into two groups in the 16S rRNA phylogeny (**Fig. 2**), while it is monophyletic and well supported in the phylogenomic tree (**Fig. 1**). This specific result has already been reported in a previous comparison between a 16S phylogeny and a phylogenomic analysis of picocyanobacteria [8] and might be due to the lower resolution of the 16S phylogeny compared to phylogenomics [43–47]. Finally, as in the phylogenomic analysis, cluster E is the closest relative to cluster D.

### Cluster D Antarctic genomes also group together in the pangenomic analysis

A total of 137,390 proteins were predicted across the 44 assemblies selected for the pangenomic analysis, which clusters these sequences into 22,886 gene clusters (GCs) (**Fig. S1**).

To better characterize cluster D and highlight the potential specificities of Antarctic taxa, we determined the GCs that are unique to the different D subgroups (**Fig. S1**). Hence, the genome assemblies of Antarctic groups D1 and D2 share 16 GCs not found in the non-Antarctic assemblies of group D3, thereby defining strictly Antarctic GCs in cluster D. Moreover, 84 GCs are present in all

D1/D2 assemblies and in some assemblies out of cluster D, but are totally absent from those of their close non-Antarctic relatives of group D3. The description of these 16 + 84 GCs is available in **Table S4**. Finally, cluster D as a whole has 33 unique GCs, while group D2/D3 assemblies share 72 unique GCs. At the group level, D1, D2 and D3 assemblies have 302, 141 and 417 unique GCs, respectively, corresponding to their accessory genomes.

### In-depth analysis of specific Antarctic gene clusters reveals potential pseudogenes in the most-closely related non-Antarctic genomes within cluster D

Focusing on the 16 + 84 GCs present in cluster D Antarctic genomes but not in the two Spanish genomes, only two of them are also enriched in terms of robust functional annotation (COG and KOfam/Pfam) with an adjusted q-value of 0.069, close to the significance threshold (GC_00000016 and GC_00002029; **Table S4**). For the other 98 GCs, their annotations are identical to those of GCs that are not specific to the Antarctic D1/D2 genomes; their functional annotations are therefore no longer specifically enriched in D1/D2.

Hence, GC_00000016 are genes that code for a transposase (*InsA*) and GC_00002029 are genes that code for a toxin-antitoxin (TA) system killer protein (*HigB*). *InsA* is present with a higher number of copies in the ULC/CS assemblies than in other assemblies (**Fig. S2**). The diversity of the *InsA* protein sequences is important with 19 different GCs. Although *HigB* shows enrichment too, it has less copies per genome than *InsA* (**Fig. S2**). Considering that HigB is a toxin protein that is part of a toxin-antitoxin (TA) system, we checked the presence of the companion antitoxin HigA and its position in the assembly. Yet, in contrast to HigB, HigA was not enriched in Antarctic genomes (**Fig. S2**).

To ensure that enrichment analyses were not affected by missed gene predictions in some assemblies, in-depth search for the genes encoding HigB and HigA was carried out with Forty-Two (**Fig. 3**) and genomic contexts were visualized with GeneSpy (**Fig. S3**). This analysis showed that HigB, when present, is almost always (37 of 38 occurrences) concomitant with HigA (**Table S5**). The alignment of the new sequences of HigB predicted by Forty-two with those of Prodigal shows that the HigB gene sequences found in *Synechococcus* sp. Cruz CV_v_12 and Tobar_12_5_mg were truncated when compared to their close relatives from Cluster D (**Fig. S4**). The genomic context of HigA-B (ORFs before HigB and after HigA) is conserved between Cruz CV_v_12 and Tobar12mg (99.5\ % ANI), but not between those and the genomes of group D2 (84.6 to 84.7% ANI) or group D1 (81.9 to 82% ANI). Instead, GeneSpy analysis revealed that HigB-A genes in group D1 and D2 are almost always bordered by, e.g., genes *HicB* (in 5 of the 7 assemblies), *HinT* (in 5 of the 7 assemblies), *Mj0435* (in 4 of the 7 assemblies) and *HEPN* (in 4 of the 7 assemblies), while HigB-A on “Cruz” and “Tobar” genomes are bordered by *WcaA* and *BcasA* genes (**Fig. S3**). Looking at the assemblies outside cluster D, Forty-Two also predicted an additional HigB sequence for *Synechococcus* sp. L2F and *Cyanobium* sp. FGCU and detailed the nature of HigB for other assemblies (e.g., undetermined or partial; Fig. 3).

**Fig. 3.**
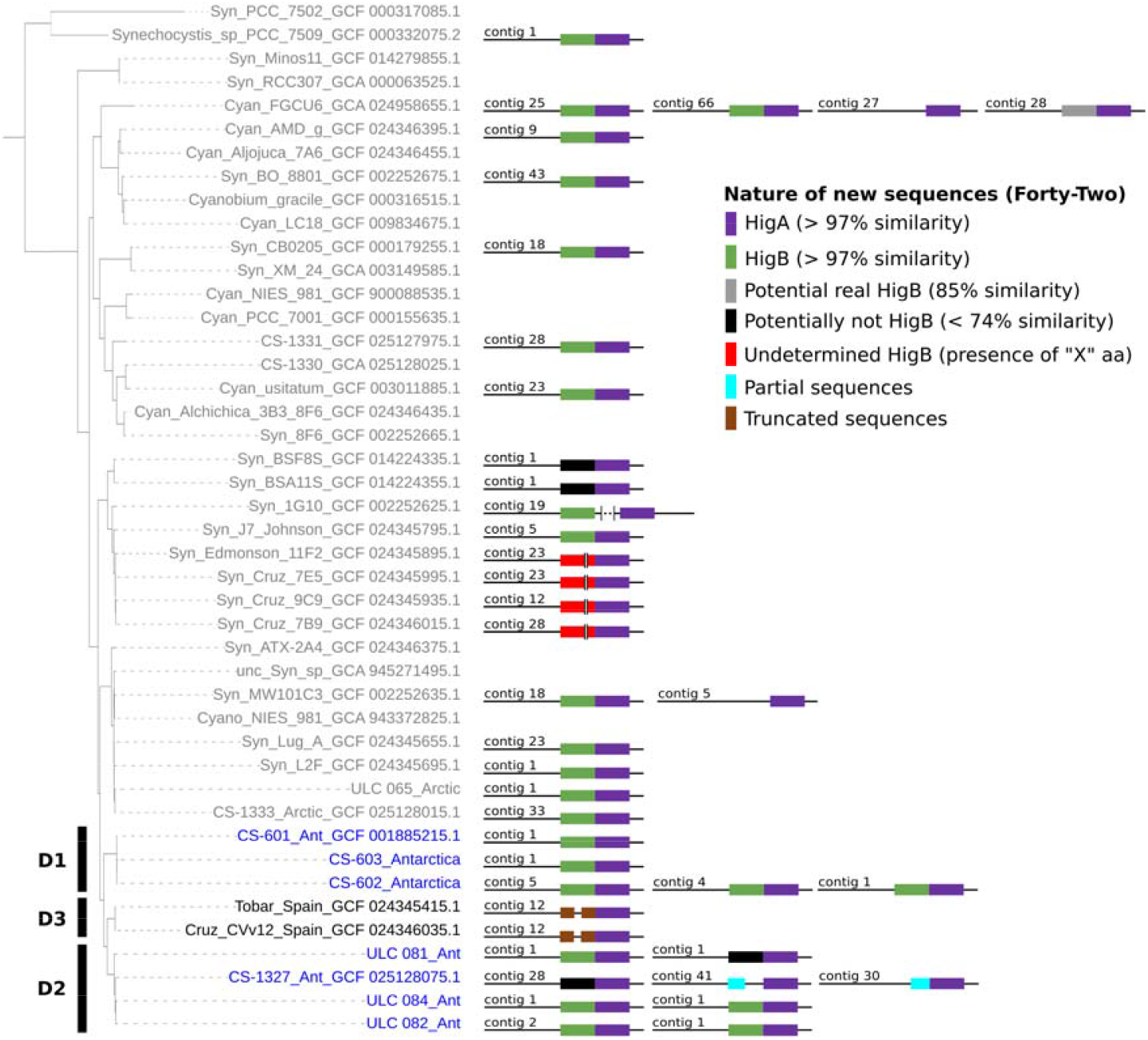
Details of the number and nature of HigA-B clusters on the 44 genomes of the pangenomic analysis, based on Forty-Two analysis.

### Functional annotations enriched in Antarctic genomes of the cluster D

Beyond the GC level, enrichment was also examined at the functional annotation level, across multiple GCs. In total, 34 and 37 functional annotations, respectively from COG and KOfam/Pfam databases, were significantly enriched (q-value < 0.05) in the D1 and D2 Antarctic genomes (T**able S6**). The intersection of these two annotations sources reveals that 19 equivalent functions (where the GCs correspond in both COG and KOfam/Pfam databases) are significantly enriched (**Fig. 4** and **Table S6**). Out of these, 21% are not predicted in detail (COG category R “General function prediction only”) and 21% are unknown (S “Function unknown); 32% of them are indirectly linked to stress or cold adaptation (**Fig. 4**).

**Fig. 4.**
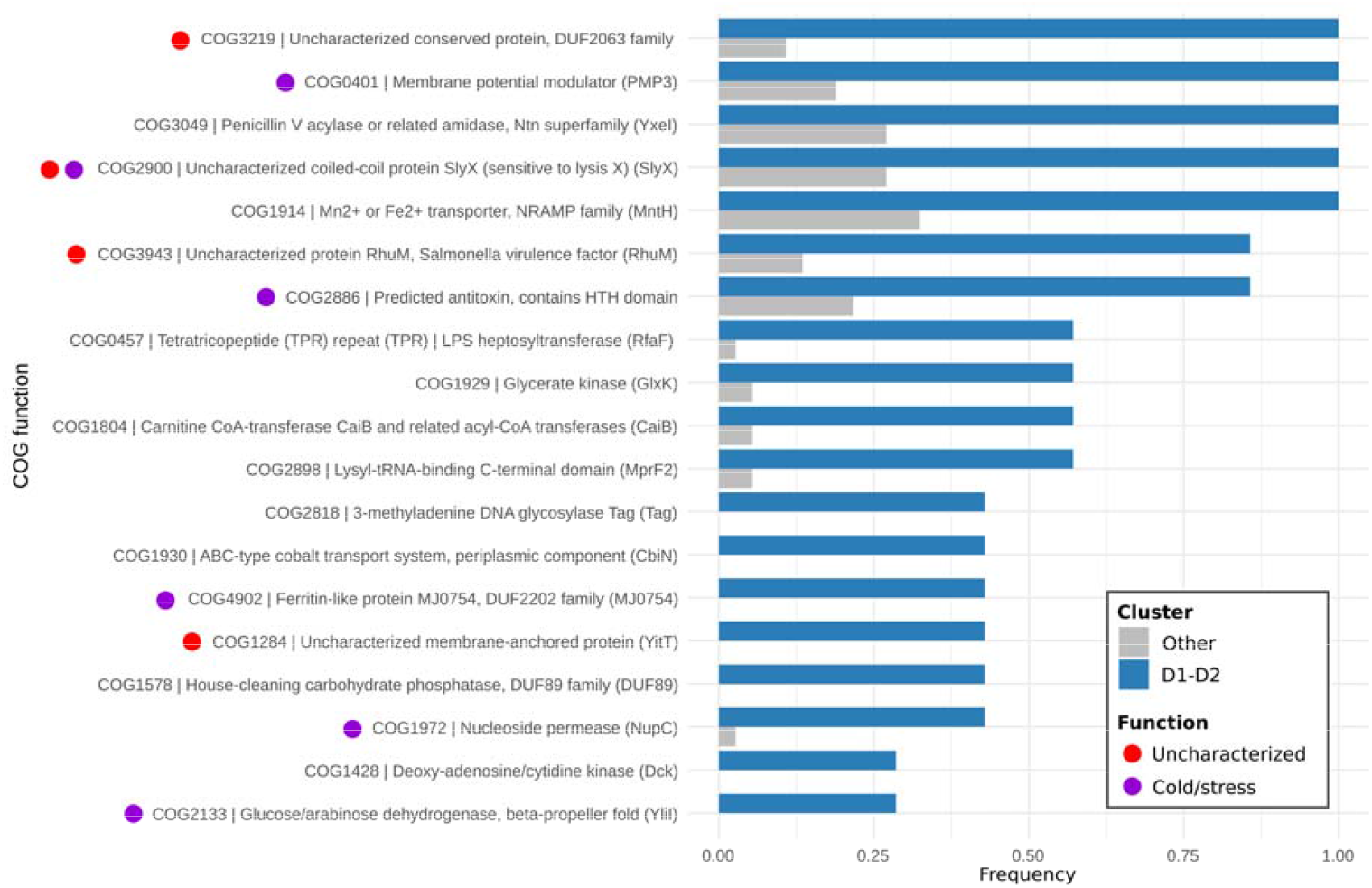
Frequency of the 19 significantly enriched functional annotations (COG ⍰ KOfam/Pfam) for the Antarctic D1-D2 group compared with the other genomes. COG functional annotations are shown; the corresponding KOfam/Pfam functional annotations are described in **Table S6**.

## 8. Discussion

### Cluster D: an Antarctic cluster as an origin for dispersal in non-Antarctic ecological niches

The subcluster 5.2 represents a particularly diversified subgroup, including marine and freshwater organisms. Among this subcluster, Antarctic genomes (e.g., CS-601, ULC084) were previously described as monophyletic in phylogenomic studies [16, 17] and 16S rRNA phylogenies [5, 16, 48], with as closest relatives two strains from the Spanish lakes La Cruz and Tobar [49]. Building on newly generated and publicly available polar genomes, our phylogenomic analysis shows that those Antarctic genomes cluster with other Antarctic taxa (CS-602, CS-603, CS-1327, ULC081, ULC082), but in a group that is intermingled with the Spanish cluster (Tobar and Cruz_CVv12) (**Fig. 1**). We designated this mostly Antarctic group within subcluster 5.2 as cluster D. Our 16S rRNA phylogenetic analysis, which specifically enriched the closest relatives of cluster D, confirms the presence of non-Antarctic strains between the two Antarctic subgroups of cluster D, even encompassing an area larger than Spain (Azores, France Pyrenees, Russia and Greenland) [8, 43–47]. Overall, our trees suggest that the ancestor of cluster D within subcluster 5.2 likely lived in Antarctic. This inference leads to the hypothesis that, in addition to being colonized several times by cyanobacteria [20], Antarctica has also been the origin of lineages that subsequently dispersed outside the continent in other ecological niches. To the best of our knowledge, apart from the hypothesis proposed for *Nodularia* in Chrismas et al. (2015), no study on cyanobacteria has yet demonstrated such an “Out-of-Antarctica” scenario.

### The specificities of genomes of the Cluster D: no clear “Antarctic signal” and a pseudogene that could be a signal of ancestry

Our pangenomic analysis of the 44 genomes determined 16 Anvi’o gene clusters (GCs) unique to the Antarctic genomes of cluster D, and 84 GCs not unique to those Antarctic genomes but strictly absent from the non-Antarctic genomes of cluster D. Due to our larger genome dataset, this number is inevitably smaller than the 901 reported in Tang et al. (2019), who compared only one Antarctic genome with three non-Antarctic ones. Moreover, Tang et al. (2019) determined the unique genes of one specific genome, whereas we determined the genes shared across a collection of genomes in the same environment.

Among these 16 + 84 GCs, we did not detect any genetic signature related to stress or cold adaptation, which could have been expected in Antarctic environments. However, two of these hundred GCs nevertheless showed functional enrichment: a transposase (InsA) and a toxin-antitoxin (TA) system killer protein (HigB). Because HigB is part of a TA system, we searched for the presence of the antitoxin *HigA* gene. HigB and HigA couples were not encoded in all genomes, but HigA was present alone, or with an altered version of HigB, in eight genomes. This is in line with other studies, in which the *HigA* gene was also found alone, a fact never reported for *HigB [50]*. When *HigA* and *HigB* genes are both present, we observed that they are encoded consecutively on the chromosome. This is also what was reported in other studies [51, 52]. The deeper analysis of HigB genes reveals the presence of truncated sequences on the two non-Antarctic (Spanish) genomes of cluster D, which could be the consequence of pseudogenization. Pseudogenization in bacteria is regarded as a process in which certain genes become disposable to the fitness of the organism due to a shift of ecological niche and hence loss of selective pressure [53, 54]. Consequently, the truncated HigB sequences found in the non-Antarctic genomes located between the two Antarctic clusters within group D phylogeny provide an additional clue for supporting an Out-of-Antarctica origin.

Beyond *InsA* and *HigB*, we identified 19 significantly enriched functional annotations in D1 and D2 Antarctic genomes (**Fig. 4**). It is no longer genes specific to genomes, but a set of genes, whose functional annotations are identical, and which, added together, show a functional enrichment. Among these enriched functional annotations, none are directly and only 32% are indirectly linked to stress or cold adaptation and may provide an ecological advantage in Antarctica. **COG0401** (membrane potential modulator PMP3/YqaE) is part of the Esi3/RCI2/PMP3 gene family that is induced in response to biotic and abiotic stresses in plants, including cold [55]. **COG1972** (nucleoside permease NupC) is part of nucleoside transporters that are involved in the cAMP/CRP-regulated incorporation of extracellular ADP-glucose enhanced under envelope stress conditions [56]. Nucleosides are essential to repair DNA damage, which is particularly important in Antarctica due to high UV radiation and oxidative stress. **COG4902** (Ferritin-like protein MJ0754) belongs to proteins that are involved in cold stress response, as reported in *Listeria monocytogenes* [57]. **COG2931** (CA2+ binding protein, RTX toxin-related) belongs to RTX toxins that use CA2+binding to modulate immune system and promote virulence [58]. CA2+ binding antifreeze protein in the Antarctic bacterium *Marinomonas primoryensis* may serve as an adhesin to transiently bind the bacteria to ice [59].

Interestingly, another component related to TA systems than the couple HigB-A described above was found to be enriched: **COG2886** (predicted antitoxin, contains HTH domain), further suggesting a potential role for TA systems in the Antarctic environment. It is possible that the absence of the antitoxin gene might lead to, e.g., the modification of transcription of genes [51, 60, 61] or the death of the bacterium [62, 63]. Moreover, another type of transposase gene than *InsA* revealed enriched: **COG2900** (Transposase IS66 family). Enrichment of transposases corroborates what Tang et al. (2019) observed in the Antarctic SynAce01 genome, which showed a high content in genes encoding diverse transposases [16]. Transpositions play a role in genomic rearrangements as well as influence gene regulation and adaptation processes that determine microevolutionary processes in cyanobacteria [64], and thus possibly in Antarctica too.

The stress and cold adaptation signal in the Antarctic genomes of group D among subcluster 5.2 might be minimized by a great diversity of locations and environmental parameters within the subcluster, which may dilute or mask lineage-specific genomic adaptations. Indeed, many strains of the subcluster 5.2 have been reported to be able to live in stressful environments, like estuaries [10, 65, 66], oligotrophic lakes [67], Black Sea [68], sub-alpine lakes [16], polar lakes (*Synechococcus* sp. CS-1330, CS-1331, CS-1332 and CS-1333), glacial lakes (*Synechococcus* sp. BO 8801, J7 Johnson, 1G10, and Lug A) and volcanic lakes (*Cyanobium* sp. Aljojuca 7A6, *Synechococcus* sp. 8F6, ATX 2A4 and *Vulcanococcus limneticus* LL). Yet, previous studies have shown that cold tolerance genes are present in most cyanobacteria, not specifically those thriving in cold environments [69, 70]. This observation is consistent with the psychrotolerance of cyanobacteria [71], which has been interpreted as evidence for a temperate origin of cold-tolerant cyanobacteria.

## Conclusion

Taken together, our phylogenomic, phylogenetic and pangenomic analyses converge toward a scenario in which the Antarctic lineage within subcluster 5.2 represents not only a diversification into polar environments, but also a potential source of dispersal toward temperate and non-polar ecological niches. The presence of truncated *HigB* gene sequences in non-Antarctic subcluster members, combined with the structure of the phylogenies provides evidence for an ancestral Antarctic origin followed by subsequent expansion outside the continent. The patchy distribution of stress-related functions suggests that cold adaptation in these lineages may reflect pre-existing psychrotolerant capacities rather than specialized Antarctic innovations, even though transposases might be of use in Antarctica.

## 9. Author statements

### 9.1 Author contributions

BD performed all analyses and prepared the Figures. VS aided with pangenomic analysis and preparation of figures, MVV aided with the gene cluster analysis, and VL contributed to the Forty-Two analysis. BD, VS, LC, DB and AW wrote the initial manuscript. All authors read and approved the final manuscript.

### 9.4 Ethical approval

Not applicable. No ethical approval was required for this study.

### 9.5 Consent for publication

Not applicable.

## 9.6 Acknowledgements

OpenAI (ChatGPT-5) was used as a support in the design of script 1, script 2 and script 3 (Figshare identifier: 10.6084/m9.figshare.30772781).

## References

1. Durieu B. Picocyanobacteria Antarctica. figshare. Dataset. 10.6084/m9.figshare.30772781.

2. Salazar VW, Tschoeke DA, Swings J, Cosenza CA, Mattoso M, et al. A new genomic taxonomy system for the Synechococcus collective. Environ Microbiol 2020;22:4557–4570.

3. Komarek J, Johansen JR, Smarda J, Strunecky O. Phylogeny and taxonomy of Synechococcuslike cyanobacteria. Fottea 2020;20:171–191.

4. Scanlan DJ, Ostrowski M, Mazard S, Dufresne A, Garczarek L, et al. Ecological Genomics of Marine Picocyanobacteria. Microbiology and Molecular Biology Reviews 2009;73:249–299.

5. Callieri C, Coci M, Corno G, Macek M, Modenutti B, et al. Phylogenetic diversity of nonmarine picocyanobacteria. FEMS Microbiol Ecol 2013;85:293–301.

6. Huber P, Diovisalvi N, Ferraro M, Metz S, Lagomarsino L, et al. Phenotypic plasticity in freshwater picocyanobacteria. Environ Microbiol 2017;19:1120–1133.

7. Doré H, Farrant GK, Guyet U, Haguait J, Humily F, et al. Evolutionary Mechanisms of Long-Term Genome Diversification Associated With Niche Partitioning in Marine Picocyanobacteria. Front Microbiol 2020;11:567431.

8. Dufresne A, Ostrowski M, Scanlan DJ, Garczarek L, Mazard S, et al. Unraveling the genomic mosaic of a ubiquitous genus of marine cyanobacteria. Genome Biol;9:R90.

9. Huang S, Wilhelm SW, Harvey HR, Taylor K, Jiao N, et al. Novel lineages of Prochlorococcus and Synechococcus in the global oceans. ISME J 2012;6:285–297.

10. Xia X, Lee P, Cheung S, Lu Y, Liu H. Discovery of Euryhaline Phycoerythrobilin-Containing Synechococcus and Its Mechanisms for Adaptation to Estuarine Environments. mSystems 2020;5:e00842–20.

11. Chen J, Li Y, Jing H, Zhang X, Xu Z, et al. Genomic and transcriptomic evidence for the diverse adaptations of Synechococcus subclusters 5.2 and 5.3 to mesoscale eddies. New Phytologist 2022;233:1828–1842.

12. Castenholz RW, Wilmotte A, Herdman M, Rippka R, Waterbury JB, et al. Phylum BX. Cyanobacteria. In: Boone DR, Castenholz RW, Garrity GM (editors). Bergey’s Manual® of Systematic Bacteriology. New York, NY: Springer New York. pp. 473–599.

13. Vincent WF, Bowman JP, Rankin LM, Mcmeekin TA. Phylogenetic diversity of picocyanobacteria in Arctic and Antarctic ecosystems. Microbial biosystems: new frontiers Proceedings of the 8th international symposium on microbial ecology 2000;1:317–322.

14. Powell LM, Bowman JP, Skerratt JH, Franzmann PD, Burton HR. Ecology of a novel Synechococcus clade occurring in dense populations in saline Antarctic lakes. Mar Ecol Prog Ser 2005;291:65–80.

15. Everroad RC, Wood AM. Phycoerythrin evolution and diversification of spectral phenotype in marine Synechococcus and related picocyanobacteria. Mol Phylogenet Evol 2012;64:381–392.

16. Tang J, Du L-M, Liang Y-M, Daroch M. Complete Genome Sequence and Comparative Analysis of Synechococcus sp. CS-601 (SynAce01), a Cold-Adapted Cyanobacterium from an Oligotrophic Antarctic Habitat. Int J Mol Sci 2019;20:152.

17. Panwar P, Williams TJ, Allen MA, Cavicchioli R. Population structure of an Antarctic aquatic cyanobacterium. Microbiome 2022;10:1–22.

18. Panwar P, Allen MA, Williams TJ, Hancock AM, Brazendale S, et al. Influence of the polar light cycle on seasonal dynamics of an Antarctic lake microbial community. 2020;1–24.

19. Durieu B, Savaglia V, Lara Y, Lambion A, Pessi IS, et al. (Sub-)Antarctic endemic cyanobacteria from benthic mats are rare and have restricted geographic distributions. Ecography 2025;1–12.

20. Chrismas N a. M, Anesio AM, Sánchez-Baracaldo P. Multiple adaptations to polar and alpine environments within cyanobacteria: a phylogenomic and Bayesian approach. Front Microbiol 2015;6:1–10.

21. Schmieder R, Edwards R. Quality control and preprocessing of metagenomic datasets. Bioinformatics 2011;27:863–864.

22. Bolger AM, Lohse M, Usadel B. Trimmomatic: a flexible trimmer for Illumina sequence data. Bioinformatics 2014;30:2114–2120.

23. De Coster W, D’Hert S, Schultz DT, Cruts M, Van Broeckhoven C. NanoPack: visualizing and processing long-read sequencing data. Bioinformatics 2018;34:2666–2669.

24. Wick RR, Judd LM, Gorrie CL, Holt KE. Unicycler: Resolving bacterial genome assemblies from short and long sequencing reads. PLoS Comput Biol 2017;13:e1005595.

25. Wood DE, Lu J, Langmead B. Improved metagenomic analysis with Kraken 2. Genome Biol 2019;20:257.

26. Marçais G, Delcher AL, Phillippy AM, Coston R, Salzberg SL, et al. MUMmer4: A fast and versatile genome alignment system. PLoS Comput Biol 2018;14:e1005944.

27. Parks DH, Imelfort M, Skennerton CT, Hugenholtz P, Tyson GW. CheckM: assessing the quality of microbial genomes recovered from isolates, single cells, and metagenomes. Genome Res 2015;25:1043–1055.

28. Cornet L, Durieu B, Baert F, D’hooge E, Colignon D, et al. The GEN-ERA toolbox: unified and reproducible workflows for research in microbial genomics. Gigascience 2023;12:1–10.

29. Stamatakis A. RAxML version 8: A tool for phylogenetic analysis and post-analysis of large phylogenies. Bioinformatics 2014;30:1312–1313.

30. Letunic I, Bork P. Interactive Tree of Life (iTOL) v6: Recent updates to the phylogenetic tree display and annotation tool. Nucleic Acids Res 2024;52:W78–W82.

31. Lagesen K, Hallin P, Rødland EA, Stærfeldt HH, Rognes T, et al. RNAmmer: Consistent and rapid annotation of ribosomal RNA genes. Nucleic Acids Res 2007;35:3100–3108.

32. Katoh K, Standley DM. MAFFT multiple sequence alignment software version 7: Improvements in performance and usability. Mol Biol Evol 2013;30:772–780.

33. Schuler GD, Epstein JA, Ohkawa H, Kans JABT-M in E. Entrez: Molecular biology database and retrieval system. Methods Enzymol 1996;266:141–162.

34. Altschul SF, Gish W, Miller W, Myers EW, Lipman DJ. Basic local alignment search tool. J Mol Biol 1990;215:403–410.

35. Eren AM, Esen OC, Quince C, Vineis JH, Morrison HG, et al. Anvi’o: An advanced analysis and visualization platformfor ‘omics data. PeerJ 2015;2015:1–29.

36. Utter DR, Borisy GG, Eren AM, Cavanaugh CM, Mark Welch JL. Metapangenomics of the oral microbiome provides insights into habitat adaptation and cultivar diversity. Genome Biol 2020;21:1–25.

37. Simion P, Philippe H, Baurain D, Jager M, Richter DJ, et al. A Large and Consistent Phylogenomic Dataset Supports Sponges as the Sister Group to All Other Animals - Forty-two Mat-Meth. Current Biology 2017;27:958–967.

38. Lupo V, Roomans C, Royen E, Ongena L, Jacquemin O, et al. Identification and characterization of archaeal pseudomurein biosynthesis genes through pangenomics. mSystems 2025;10:1–24.

39. Sievers F, Wilm A, Dineen D, Gibson TJ, Karplus K, et al. Fast, scalable generation of highquality protein multiple sequence alignments using Clustal Omega. Mol Syst Biol 2011;7:539.

40. Guindon S, Delsuc F, Dufayard J-F, Gascuel O. Estimating maximum likelihood phylogenies with PhyML. Methods Mol Biol 2009;537:113–137.

41. Guindon S, Dufayard JF, Lefort V, Anisimova M, Hordijk W, et al. New algorithms and methods to estimate maximum-likelihood phylogenies: Assessing the performance of PhyML 3.0. Syst Biol 2010;59:307–321.

42. Garcia PS, Jauffrit F, Grangeasse C, Brochier-Armanet C. GeneSpy, a user-friendly and flexible genomic context visualizer. Bioinformatics 2019;35:329–331.

43. Gontcharov AA, Marin B, Melkonian M. Are Combined Analyses Better Than Single Gene Phylogenies? A Case Study Using SSU rDNA and rbcL Sequence Comparisons in the Zygnematophyceae (Streptophyta). Mol Biol Evol 2004;21:612–624.

44. Dessimoz C, Gil M. Phylogenetic assessment of alignments reveals neglected tree signal in gaps. Genome Biol 2010;11:R37.

45. Lunter G, Rocco A, Mimouni N, Heger A, Caldeira A, et al. Uncertainty in homology inferences: Assessing and improving genomic sequence alignment. Genome Res 2008;18:298–309.

46. Wong KM, Suchard MA, Huelsenbeck JP. Alignment Uncertainty and Genomic Analysis. Science (1979) 2008;319:473–476.

47. Mareš J. Multilocus and SSU rRNA gene phylogenetic analyses of available cyanobacterial genomes, and their relation to the current taxonomic system. Hydrobiologia 2018;811:19–34.

48. Everroad RC, Woebken D, Singer SW, Burow LC, Kyrpides N, et al. Draft Genome Sequence of an Oscillatorian Cyanobacterium, Strain ESFC-1. Genome Announc 2013;1:13–14.

49. Cabello-Yeves PJ, Callieri C, Picazo A, Schallenberg L, Huber P, et al. Elucidating the picocyanobacteria salinity divide through ecogenomics of new freshwater isolates. BMC Biol 2022;20:1–24.

50. Budde PP, Davis BM, Yuan J, Waldor MK. Characterization of a higBA toxin-antitoxin locus in Vibrio cholerae. J Bacteriol 2007;189:491–500.

51. Fraikin N, Goormaghtigh F, Van Melderen L. Type II Toxin-Antitoxin Systems: Evolution and Revolutions. J Bacteriol 2020;202:e00763–19.

52. Deter HS, Jensen R V., Mather WH, Butzin NC. Mechanisms for differential protein production in toxin–antitoxin systems. Toxins (Basel) 2017;9:1–13.

53. Kuo CH, Ochman H. The extinction dynamics of bacterial pseudogenes. PLoS Genet 2020;6(8): e1001050.

54. Goodhead I, Darby AC. Taking the pseudo out of pseudogenes. Curr Opin Microbiol 2015;23:102–109.

55. Brunetti SC, Arseneault MKM, Gulick PJ. Characterization of the Esi3/RCI2/PMP3 gene family in the Triticeae. BMC Genomics 2018;19:898.

56. Almagro G, Viale AM, Montero M, Muñoz FJ, Baroja-Fernández E, et al. A cAMP/CRP-controlled mechanism for the incorporation of extracellular ADP-glucose in Escherichia coli involving NupC and NupG nucleoside transporters. Sci Rep 2018;8:15509.

57. Miladi H, Soukri A, Bakhrouf A, Ammar E. Expression of ferritin-like protein in Listeria monocytogenes after cold and freezing stress. Folia Microbiol (Praha) 2012;57:551–556.

58. Coote JG. Structural and functional relationships among the RTX toxin determinants of Gramnegative bacteria. FEMS Microbiol Lett 1992;88:137–162.

59. Guo S, Garnham CP, Whitney JC, Graham LA, Davies PL. Re-Evaluation of a Bacterial Antifreeze Protein as an Adhesin with Ice-Binding Activity. PLoS One 2012;7:e48805.

60. Liu Y, Gao Z, Liu G, Geng Z, Dong Y, et al. Structural Insights Into the Transcriptional Regulation of HigBA Toxin–Antitoxin System by Antitoxin HigA in Pseudomonas aeruginosa. Front Microbiol 2020;10:1–13.

61. Qin Q, Muhammad K, R. IJ. The higBA-Type Toxin-Antitoxin System in IncC Plasmids Is a Mobilizable Ciprofloxacin-Inducible System. mSphere 2021;6:10.1128/msphere.00424-21.

62. Singh G, Yadav M, Ghosh C, Rathore JS. Bacterial toxin-antitoxin modules: classification, functions, and association with persistence. Curr Res Microb Sci 2021;2:100047.

63. Hurley JM, Woychik NA. Bacterial Toxin HigB Associates with Ribosomes and Mediates Translation-dependent mRNA Cleavage at A-rich Sites*. Journal of Biological Chemistry 2009;284:18605–18613.

64. Mikheeva LE, Karbysheva EA, Shestakov S V. The role of mobile genetic elements in the evolution of cyanobacteria. Russ J Genet Appl Res 2013;3:91–101.

65. Zhu X, Li Q, Yin C, Fang X, Xu X. Role of spermidine in overwintering of cyanobacteria. J Bacteriol 2015;197:2325–2334.

66. Fucich D, Marsan D, Sosa A, Chen F. Complete Genome Sequence of Subcluster 5.2 Synechococcus sp. Strain CB0101, Isolated from the Chesapeake Bay. Microbiol Resour Announc 2019;8:e00484–19.

67. Cabello-Yeves PJ, Picazo A, Camacho A, Callieri C, Rosselli R, et al. Ecological and genomic features of two widespread freshwater picocyanobacteria. Environ Microbiol 2018;20:3757– 3771.

68. Di Cesare A, Dzhembekova N, Cabello-Yeves PJ, Eckert EM, Slabakova V, et al. Genomic Comparison and Spatial Distribution of Different Synechococcus Phylotypes in the Black Sea. Front Microbiol 2020;11:1979.

69. Chrismas NAM, Barker G, Anesio AM, Sánchez-baracaldo P. Genomic mechanisms for cold tolerance and production of exopolysaccharides in the Arctic cyanobacterium Phormidesmis priestleyi BC1401. BMC Genomics 2016;1–14.

70. Lumian J, Sumner DY, Grettenberger CL, Jungblut AD, Irber L, et al. Biogeographic distribution of five Antarctic cyanobacteria using large-scale k-mer searching with sourmash branchwater. Front Microbiol 2024;15:1–10.

71. Tang EPY, Tremblay R, Vincent WF. Cyanobacterial dominance of polar freshwater ecosystems: are high-latitude mat-formers adapted to low temperature? 1. J Phycol 1997;33:171–181.

