## Supplementary figures for "Out of Antarctica: new insights into Antarctic Subcluster 5.2 picocyanobacteria based on high-quality genomes"

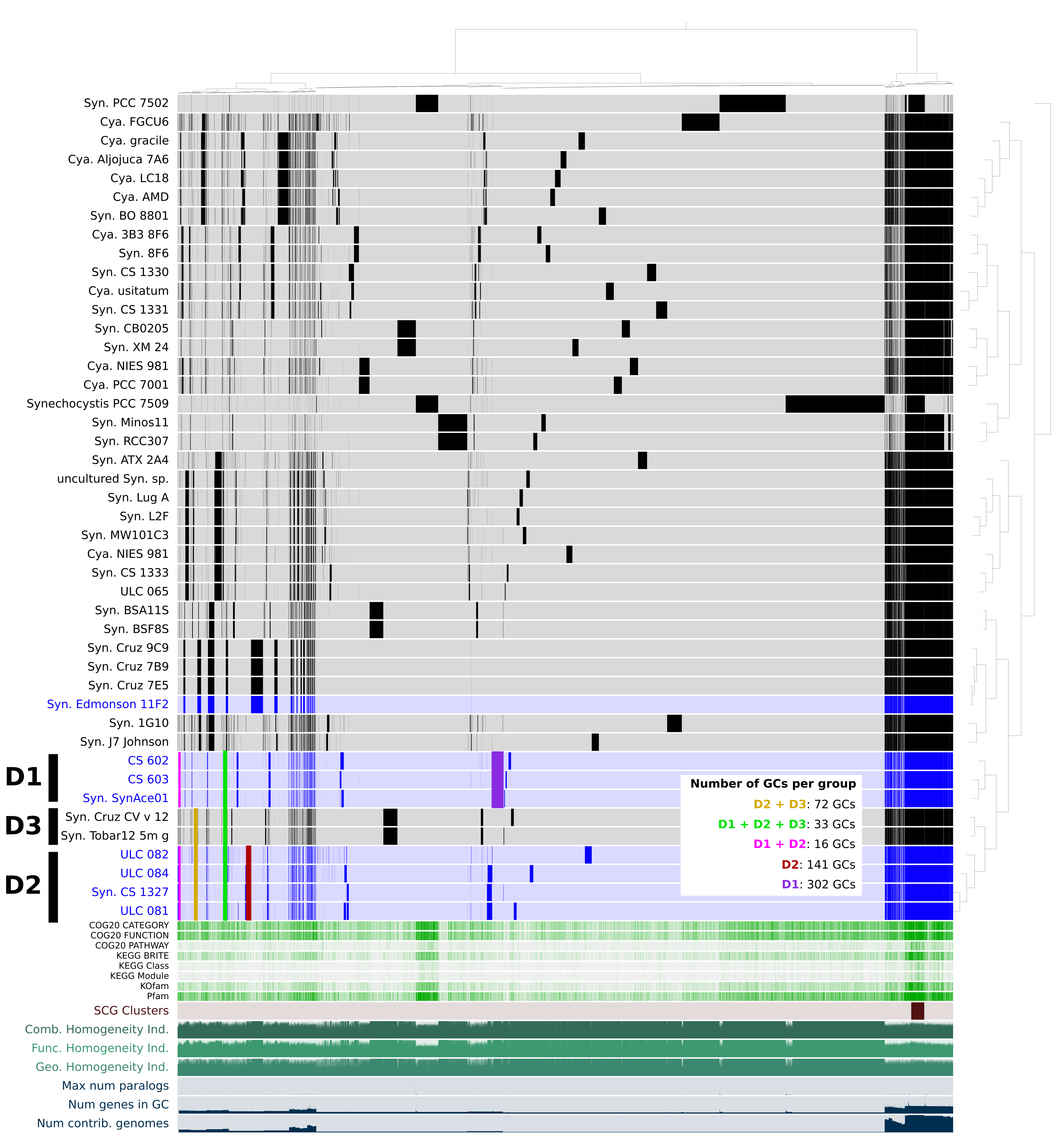

[Fig. S1](https://mseduculiegebe-my.sharepoint.com/:i:/r/personal/benoit_durieu_doct_uliege_be/Documents/Chap4_paper_Picocyano/Figures/Figure_S1_genomes_44_PAN_minbit_0_5_mcl_8_25022_modif_2.png?csf=1&web=1&e=sk4fz8). Pangenomic analysis of the 44 cyanobacterial assemblies (minbit = 0.5, mcl inflation = 8) revealing 5 groups of GCs linked to Antarctic genomes of cluster D (groups ’D2 + D3‘, ‘D1 + D2 + D3’, ’D1 + D2’, ‘D2’ and ‘D1‘).

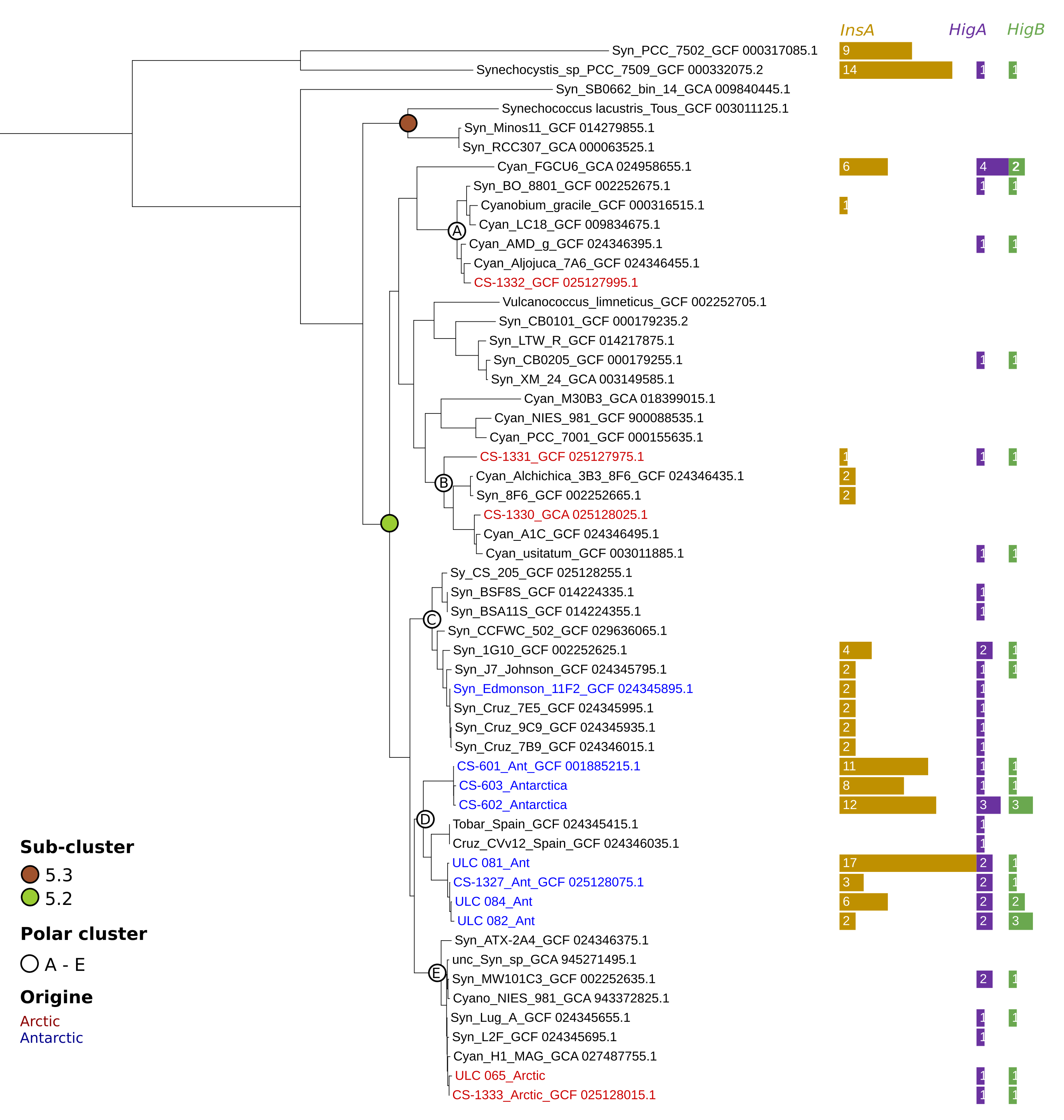

[Fig. S2](https://mseduculiegebe-my.sharepoint.com/:i:/r/personal/benoit_durieu_doct_uliege_be/Documents/Chap4_paper_Picocyano/Figures/Figure_S2_Tree_55_InsA_HigA-B_update_110325.png?csf=1&web=1&e=nbBijp). Multilocus maximum likelihood phylogeny with 1000 rapid bootstraps under the PROTGAMMALGF model of 55 cyanobacterial assemblies inferred from a supermatrix of 476 genes totalizing 86,369 unambiguously aligned amino acid positions and 0.21% of missing characters. Number of copies of InsA (gold), HigA (purple) and HigB (green) genes predicted by Anvi’o workflow are represented by horizontal barplots.

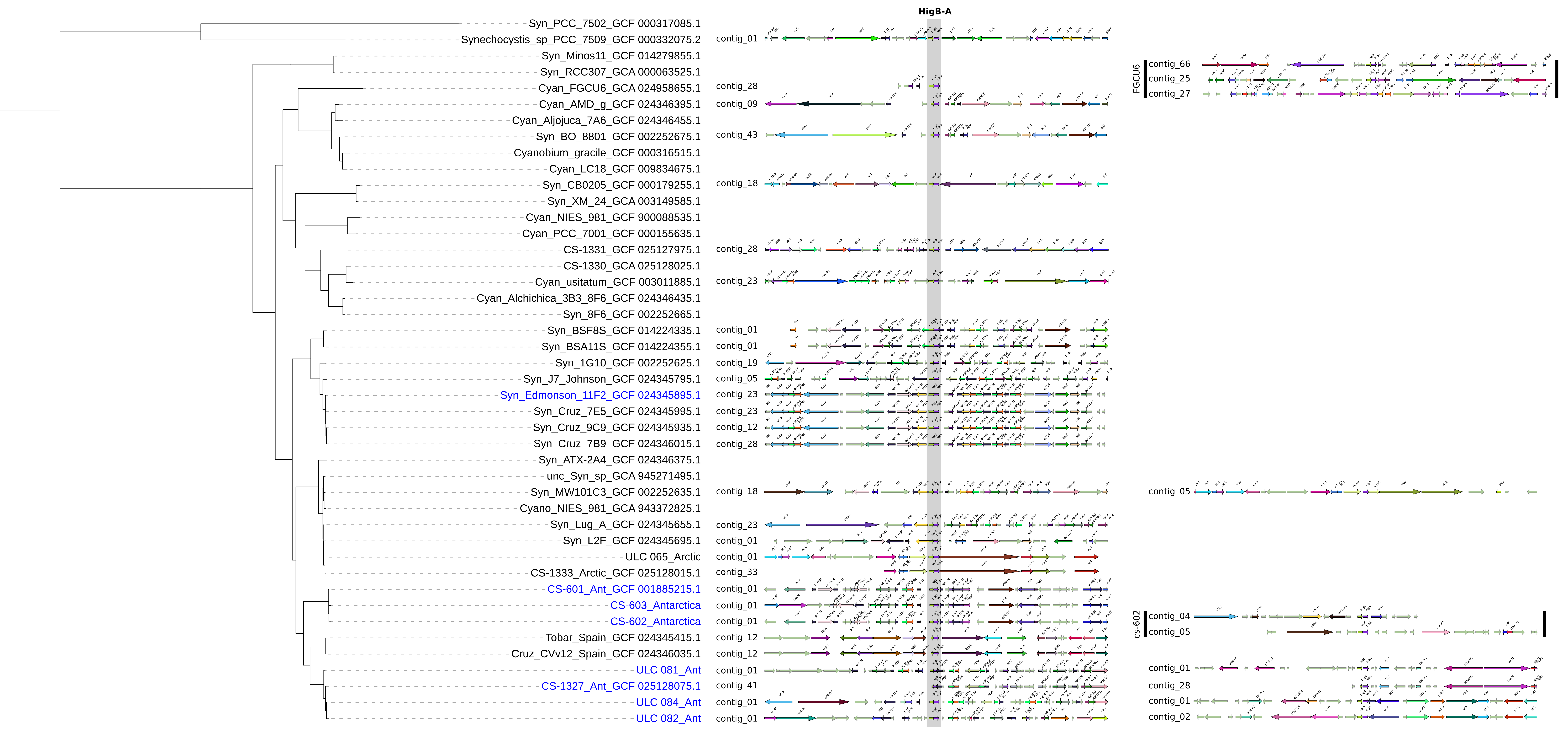

[Fig. S3](https://mseduculiegebe-my.sharepoint.com/:i:/r/personal/benoit_durieu_doct_uliege_be/Documents/Chap4_paper_Picocyano/Figures/Phylo_GeneSpy_HigB-A.png?csf=1&web=1&e=08E107). Genomic contexts of HigA and HigB protein sequences in the 44 assemblies.

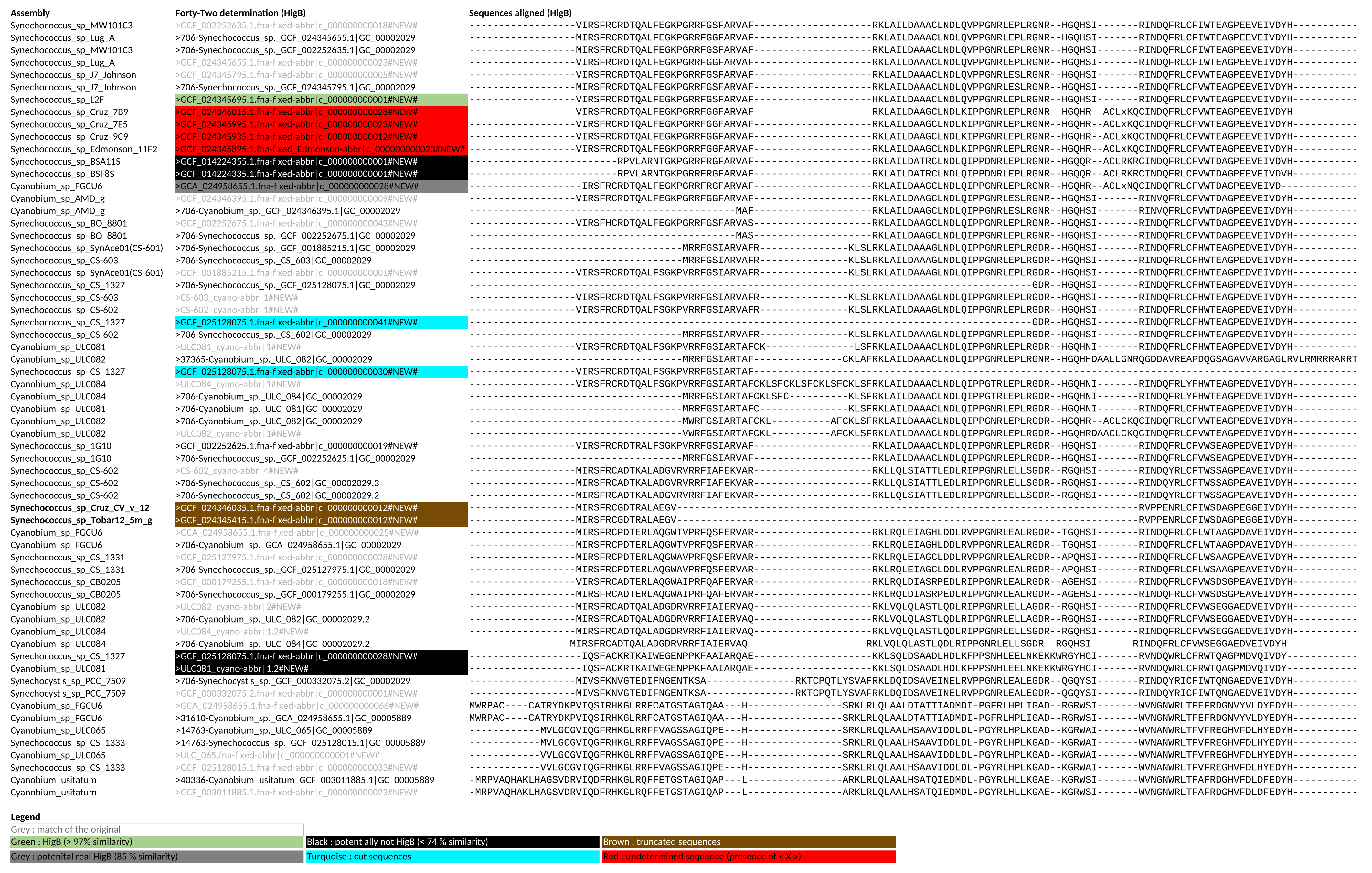

[Fig. S4](https://mseduculiegebe-my.sharepoint.com/:i:/r/personal/benoit_durieu_doct_uliege_be/Documents/Chap4_paper_Picocyano/Figures/Tab_S9_HigA-B_Forty-two_Alignement.png?csf=1&web=1&e=EhrIBY). Alignment of the HigB protein sequences predicted by Anvi’o and Forty-Two (colored).
